# Mechanism of Phosphatidylserine Lipid Scrambling by Human SERINC3, an HIV-1 Restriction Factor

**DOI:** 10.1101/2025.11.13.688349

**Authors:** Puja Banerjee, Mark Yeager, Gregory A. Voth

## Abstract

The HIV-1 restriction factor, hSERINC3, functions as a lipid scramblase, translocating lipids across the bilayer in reconstituted proteoliposomes and the viral envelope. Phosphatidylserine(PS) scrambling and exposure at the outer leaflet are recognized to play important roles in several biological processes. To understand the mechanistic basis for hSERINC3-mediated PS lipid scrambling at atomistic resolution, we implemented the transition-tempered metadynamics (TTMetaD) enhanced sampling method. Our simulations sampled close-to-open hSERINC3 conformational transition during PS scrambling and demonstrated that while other non-ATP-dependent lipid transporters with similar architecture transport lipid following a “trap-and-flip” mechanism, hSERINC3 adopts a “credit card” mechanism of lipid scrambling and does not follow the classical “alternating access” mechanism. Notably, we observe unfolding of the H8 NTD, consistent with the cryo-EM density map of WT-hSERINC3, mediates PS scrambling. A cluster of hydrophilic residues in the hSERINC3 central cavity, forming central gates and interacting with the PS headgroup, stabilizes the intermediate state of inner-groove scrambling and is also observed in the AlphaFold2 model of hSERINC5 that exhibits the highest viral restriction activity. Surprisingly, our simulations reveal distinct pathways for lipid translocation and pathway-dependent alterations of hSERINC3 central cavity, providing direct evidence for a non-canonical, closed-state out-of-groove PS scrambling in a complex membrane environment.

## Introduction

A fundamental feature of the host cell plasma membrane (PM) is the asymmetric distribution of lipids between the inner and outer leaflets^1^. HIV-1 assembles at the plasma membrane, and is released by fission, thereby maintaining lipid asymmetry of the plasma membrane in the viral envelope. Lipid asymmetry of any membrane is established and typically maintained by ATP-dependent, unidirectional lipid transporters, such as flippases and floppases^2^. In contrast, scramblases enable ATP-independent, bidirectional lipid transport that disrupts normal membrane asymmetry ^3, 4^. Phosphatidylserine (PS) lipid scrambling and exposure at the outer leaflet or cell surface play a key role in orchestrating cell-cell interactions, signaling, and biochemical reactions involved in a wide range of physiological processes, including phagocytosis of apoptotic cells, platelet activation, and bone mineralization^5, 6^.

Cellular PMs and HIV-1 envelope membranes are composed of various kinds of phospholipids with monounsaturated or polyunsaturated tails. The inner, cytoplasmic leaflet of these asymmetric membranes contains highly negatively charged phospholipids such as phosphatidylserine (PS) and phosphatidylinositol (PI), and overall a greater number of unsaturated (loosely packed) lipids than the outer leaflet, which play diverse roles in viral assembly and maturation phases^7–10^. The outer leaflet primarily contains phosphatidylcholine (PC), sphingolipid, and cholesterol to build a highly ordered and rigid layer with low permeability. Asymmetry in the lipid distribution, lipid unsaturation, and differential packing of lipids generate curvature and alter the physical properties of the membrane such as bilayer thickness^1^. Loss of asymmetry eliminates differential stress and alters membrane physical properties. The role of scramblases in relaxing membrane asymmetry is widely accepted, and several scramblases have been identified and characterized ^6, 11, 12^. Nevertheless, the mechanism of lipid scrambling mediated by the protein-lipid interactions is not fully understood.

In addition to experimental results, several computational approaches have been used to gain insight into protein-lipid interactions that enable lipid transport, such as molecular dynamics (MD) simulations using atomistic or coarse-grained (CG) forcefields, which can be combined with advanced simulation methods such as the Markov state modeling, targeted MD, as well as MD simulation with an external electric field ^13–21^. Depending on the protein-lipid interactions, two different models have been proposed, the “trap-and-flip” and the “credit-card” mechanisms. The “credit card” mechanism of protein-mediated lipid flipping was first proposed by Menon and Pomorski, which was recapitulated by simulations^22,13^, suggesting that the headgroup of a translocating phospholipid traverses through a hydrophilic groove of the protein, while the hydrophobic, aliphatic lipid tails interact with the membrane. In contrast, the “trap-and-flip” mechanism proposes that lipid molecules being translocated remain completely enclosed within a central cavity of the protein. Advances in structure determination and computer simulation have revealed that both of these mechanisms can follow or be distinct from the classical “alternating-access model”, which describes protein conformational change between three states (inward-open, occluded, and outward-open), to accommodate the lipid and facilitate the transport. The characterization of lipid transport models for different classes of membrane transporters and the atomic-scale determination of the lipid translocating pathways, crucial protein-lipid interactions, and the energy barrier along those pathways remain an outstanding scientific challenge.

The serine incorporator (SERINC) family of proteins are integral, transmembrane proteins that are believed to be associated with the biosynthesis of serine-derived lipids, PS, and sphingolipids. Two isoforms, human SERINC3 (hSERINC3) and hSERINC5, display potent restriction activity against HIV-1 and some other enveloped viruses ^23–25^. For HIV-1, restriction activity is counteracted by the viral protein, Nef ^23^. Interestingly, hSERINC2 lacks restriction activity. In the previous studies by Leonhardt *et al.* ^26^, using a fluorescent proteoliposome assay, it was demonstrated that hSERINC3 flips PS, PC, and phosphatidylethanolamine (PE). Such nonspecific lipid flipping is a hallmark of scramblases. Experiments in HIV-1 particles^26^ demonstrated lipid-flipping activity of PS and the consequent disruption of membrane asymmetry are strongly correlated with changes in restriction activity and the conformation of the envelope trimer (Env). A separate study demonstrated that SERINCs modulate viral membrane properties, such as lipid chain order, rigidity, line tension, and lateral pressure^27^. To understand the potential local or large-scale biophysical alterations of the membrane properties induced by SERINCs, and in turn the structural change of other membrane proteins^28, 29^, such as Env, we need to first understand the SERINC-membrane interactions in atomistic detail. Here, we considered the cryo-EM-derived structure of hSERINC3^26^ to understand the specific protein-membrane interactions, while focusing on the lipid scrambling activity by hSERINC3.

To understand the mechanism of hSERINC3-mediated PS lipid scrambling, we have performed atomistic MD simulations with a unique enhanced sampling method. Here we report the computation of the free-energy surface for multi-pathway protein-mediated lipid scrambling in a complex asymmetric membrane at atomistic resolution employing the transition-tempered metadynamics (TTMetaD) method^30^. For the biomolecular processes with known initial and final states, this is a unique modification of the well-tempered metadynamics method ^31, 32^, a technique widely recognized as a powerful tool for probing complex biomolecular interactions ^33–36^. In TTMetaD, the biasing scheme does not presume beforehand the pathways of lipid transport, and it allows the exploration of the free energy hypersurface of hSERINC3-PS headgroup interactions during lipid translocation through protein grooves, avoiding other competitive processes such as lipid lateral diffusion and lipid desorption from the membrane. TTMetaD also rapidly converges the free energy surface for the chosen collective variables (CVs).

An open conformation of the membrane transporter has been essential for simulations that investigate lipid permeation events^13, 16, 19–21^. Coarse-grained simulations using the MARTINI forcefield are known to be limited in sampling correlated protein close-to-open conformational change during lipid scrambling and capturing the energetics of lipid translocation^15, 37^. Our combined MD and TTMetaD methodology enabled the study of lipid scrambling mechanisms starting with a non-open conformation of hSERINC3 as it allowed close-to-open transitions of the protein groove in an unsupervised way. To the best of our knowledge, our simulations provide the first atomistic description of multi-pathway protein-mediated lipid scrambling dynamics, allowing spontaneous closed-to-open protein conformational transitions during lipid translocation. By investigating conformational changes of hSERINC3 and hSERINC3-PS specific interactions during PS lipid scrambling, the results presented in this work reveal important molecular-level mechanistic details of hSERINC3-mediated PS lipid scrambling. Additionally, this unique mechanism expands the repertoire of protein-mediated lipid scrambling, which may be relevant for the design of novel therapeutics to modulate the function of such scramblases.

## Results

### PS lipids bind hSERINC3 close to its cytoplasmic entrance in the asymmetric membrane

To investigate how PS interacts with hSERINC3, we performed long-timescale unbiased all-atom molecular dynamics (AAMD) simulations of the cryoEM density map-derived hSERINC3 structure (PDB ID: 7RU6) embedded in an asymmetric membrane model resembling the HIV-1 envelope. The cytoplasmic, i.e., intracellular leaflet (ICL), unlike the extracellular leaflet (ECL), contains highly negatively charged lipids, including phosphatidylinositol 4,5-bisphosphate (PIP2) and PS, which interact with basic residues on the cytoplasmic face of hSERINC3. Long unbiased MD simulations maintained the conformational state of hSERINC3 with a narrow transmembrane aqueous channel, displayed no phospholipid flipping, and are referred to as (-)PS-scrambling simulations throughout the manuscript. Multiple replicas of these simulations revealed the specific PS-binding sites on the cytoplasmic side of the protein. The cytoplasmic cavity, formed by H5, H7, H8, and the C-terminal end of the H4 crossmember helix, was identified as a key interaction region (**Figure 1**). The PS headgroup density map, computed from concatenated unbiased MD trajectories, highlights high-density domains (green in (**Figure 1B**)) near the cytoplasmic entrance. The basic residues of H4, H5, and H8 at the cytoplasmic side (ARG193, ARG200, and ARG354) bind the negatively charged PS headgroups.

**Figure 1.**
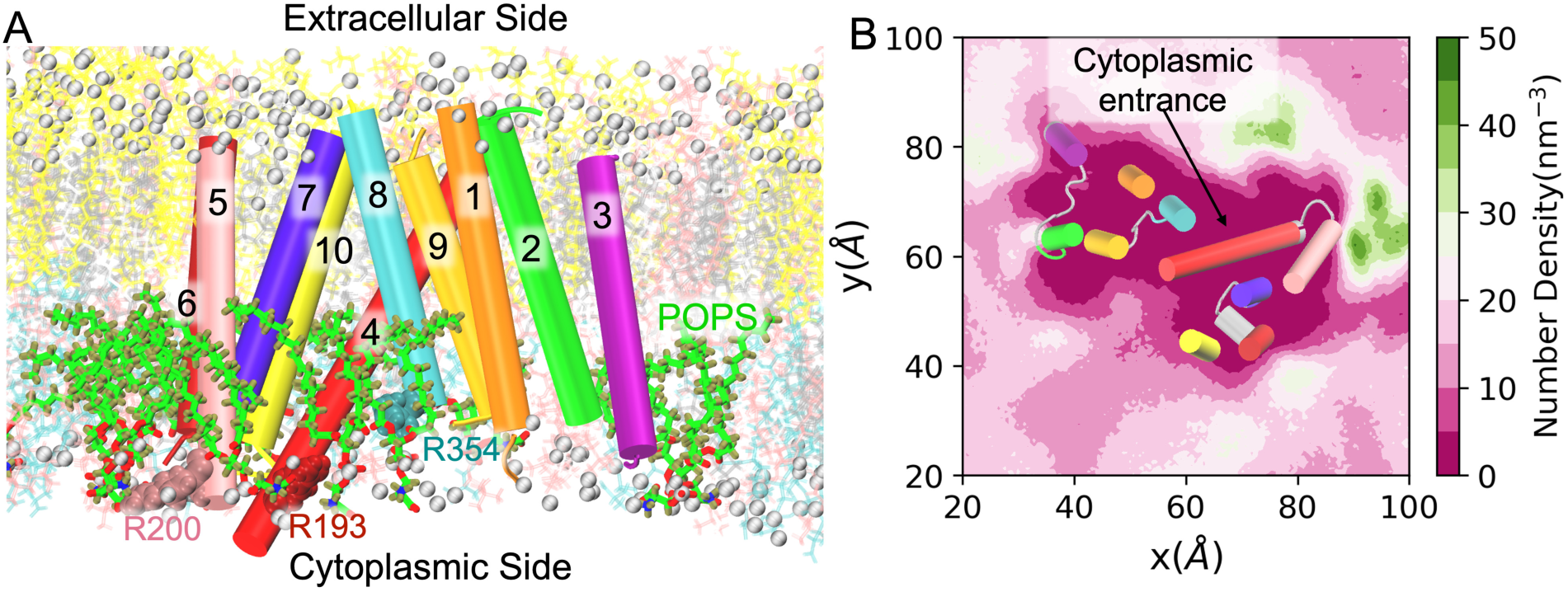
: Unbiased AAMD simulations reveal PS enrichment near the cytoplasmic entrance of hSERINC3. hSERINC3 is comprised of two α-helical bundles (H5, H6, H7, H10 and H1, H2, H3 H9) connected by a highly tilted 40-residue H4 crossmember helix. H8 packs against H4. (A) Simulation snapshot of a side view of hSERINC3, embedded in an asymmetric bilayer. PS lipids (green tails and red headgroup atoms) interacting with basic residues on the cytoplasmic side of the membrane. ARG residues of H4, H5, and H8 are displayed in the colors of their corresponding α-helices. (B) Density map of PS lipid headgroups around hSERINC3 shows a higher number density (green color) near the cytoplasmic entrance. Positions of cytoplasmic leaflet-spanning α-helices (as viewed from the extracellular side) are shown for reference.

### Free-energy surface of multi-pathway hSERINC3-mediated PS scrambling

We computed the free energy surface (FES) for hSERINC3-PS headgroup interactions during the scrambling process as a function of appropriate collective variables (CVs) using the TTMetaD method (see ***Methods*** section for details). TTMetaD trajectories are hereafter referred to as (+)PS-scrambling in the Results and Discussion sections. The chosen collective variables (CVs) track the movement of lipid headgroups between two leaflets (z-distance) and the interactions of lipid headgroups with protein residues in/out of the central cavity (y-distance). The FES (**Figure 2A**) reveals two distinct intermediate states (IS1, IS2), corresponding to separate scrambling pathways. In pathway 1, the PS headgroup traverses through the inner groove or central cavity of hSERINC3 (y-distance ∼ 0 Å; **Figure 2D**). In pathway 2, the PS headgroup interacts with the residues at the protein surface (y-distance ∼ 15 Å; **Figure 2E**). Although there can be many possible pathways on this complex FES connecting stable states of PS at ICL or ECL, here we have shown two probable low-barrier pathways through IS1 and IS2. The higher free energy domains (>18 kcal/mol; red color in **Figure 2A**) correspond to the infeasible PS lipid scrambling domains, such as the tightly packed interiors of the two helical bundles, H1-H2-H3-H9, and H5-H6-H7-H10.

**Figure 2:**
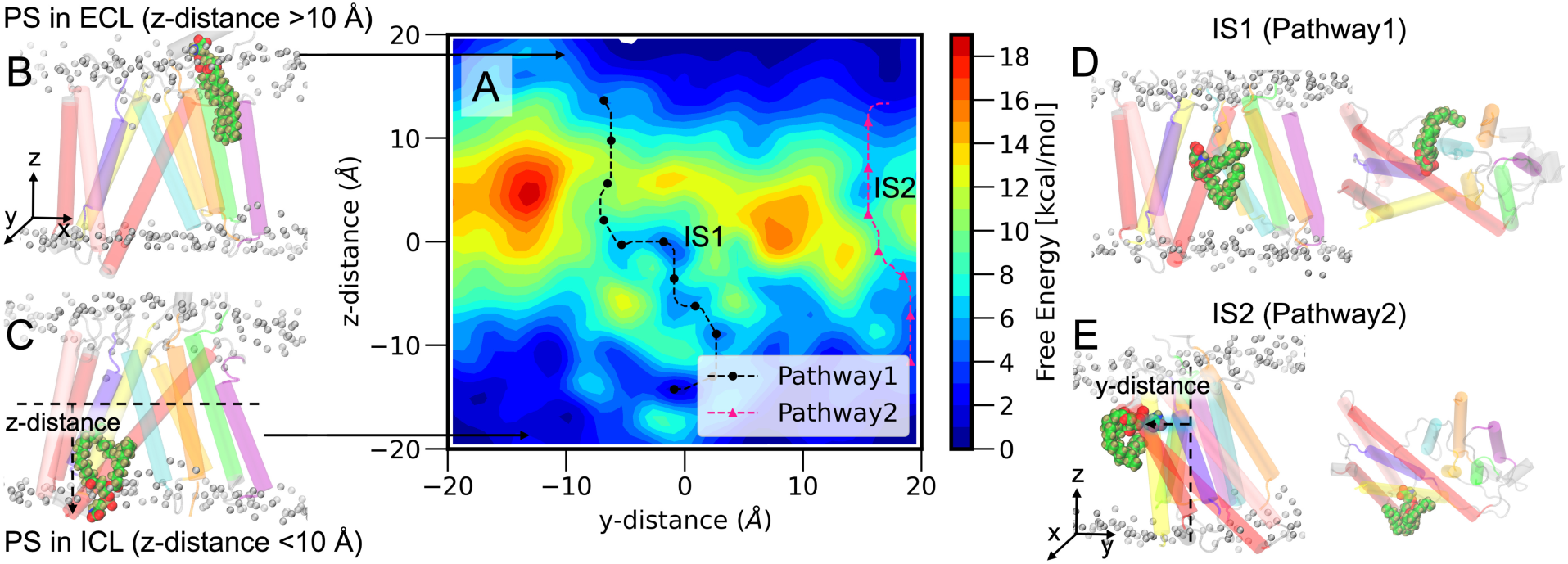
Computational analysis reveals pathways of PS lipid scrambling through hSERINC3. A. Free energy surface as a function of two selected CVs: z-distance, representing the flipping coordinate, and y-distance, representing insertion of the lipid headgroups into the central cavity of hSERINC3. (B,C) Snapshots of stable basins with the PS headgroup located in the extracellular leaflet (ECL) or intracellular leaflet (ICL). (D,E) Snapshots of two intermediate states (IS1, IS2) along distinct scrambling pathways: the PS headgroup is at the inner groove for IS1 and at the surface of hSERINC3 at IS2.

### Helix conformational changes, rearrangements, and hydrophobic gates of hSERINC3 mediate PS lipid scrambling

To investigate conformational changes in hSERINC3 during PS translocation, we compared unbiased ((-)PS-scrambling) and TTMetaD ((+)PS-scrambling) MD simulation trajectories (***Figure 3***). Under (+)PS-scrambling conditions, the cytoplasmic ends of the H8 and H10 helices increased their separation by ∼10 Å (***Figure 3*D**). Notably, the H8 helix, positioned near the inner groove, undergoes partial unfolding near the narrow extracellular vestibule in the (+)PS-scrambling simulations. The helicity parameter fluctuates around 16.0 (maximum value for 21 residues, ASP334-ARG354) under the (-)PS-scrambling condition, which makes a transition to ∼12.5 by partial unfolding of H8 residues ASP334-ASN337 under the (+)PS-scrambling condition (***Figure 3*B-D**, **Figure S1** A, D). This behavior aligns well with the WT hSERINC3 cryo-EM density map, which shows poorly defined density for H8 in the extracellular leaflet (ECL)^26^ (***Figure 3*A**). During (+)PS-scrambling simulations, large-scale rearrangements of transmembrane helices render the central cavity accessible to the surrounding membrane (***Figure 3*E**, **Figure S1** B,E). Non-rigid motions of helical bundles H1–H3, H9, and H5–H7, H10 were detected and quantified using time-structure independent component analysis^38^ (tICA) (**Figure S7**), in contrast to predictions from cryo-EM data ^26^. Additional conformational variability in local twisting and bending was observed for H4, consistent with experimental results (**Figure S6**).

**Figure 3:**
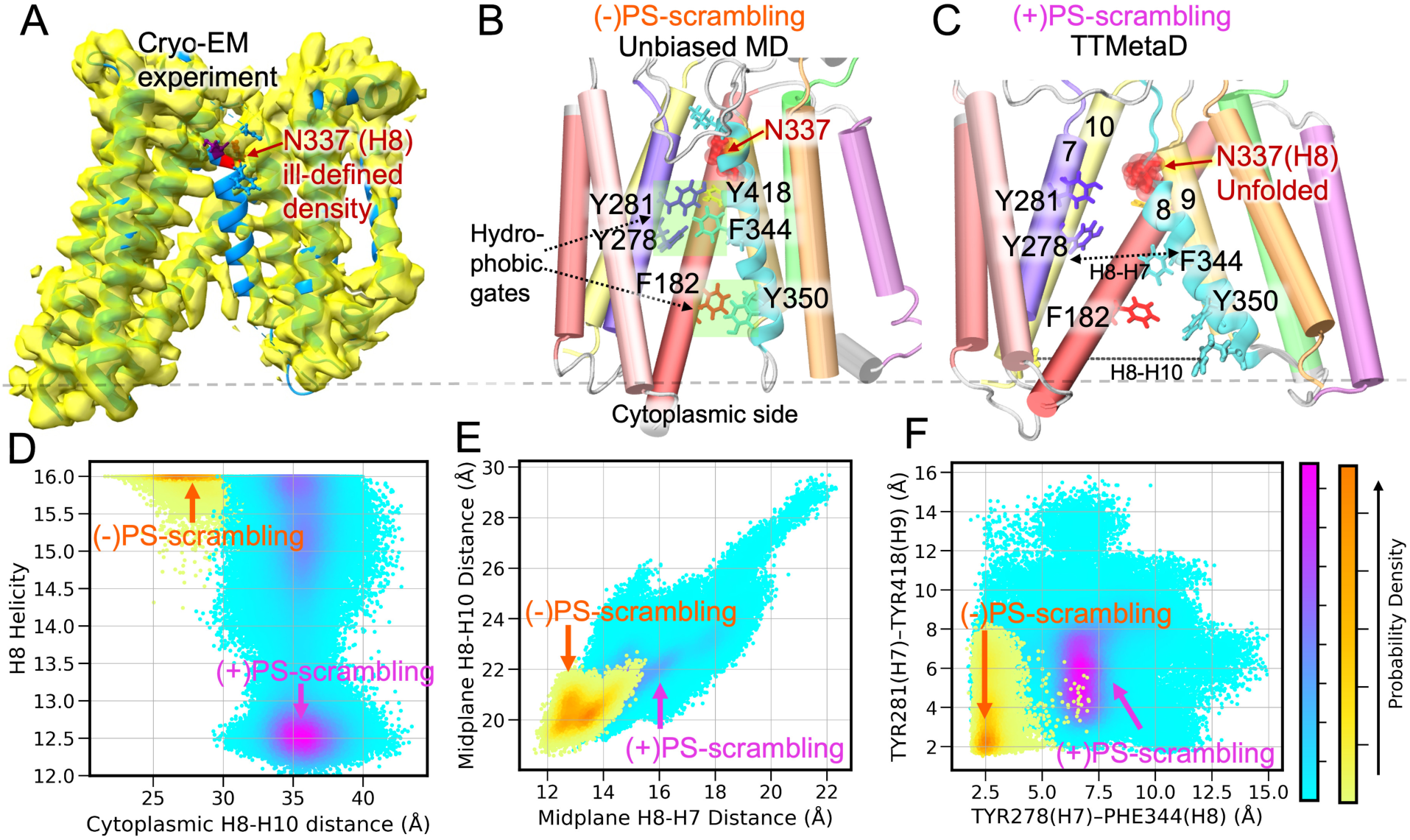
Conformational changes in hSERINC3 associated with PS scrambling activity. (A) The WT hSERINC3 cryo-EM model and map showing poorly defined density for H8. ASN337 from the disordered domain of H8 is highlighted in red. (B) The helical conformation of H8 (cyan) of the initial structure (PDB ID:7RU6) is maintained in the unbiased MD simulations of hSERINC3 for the (-)PS-scrambling condition. (C) TTMetaD simulations under the (+)PS-scrambling condition sample partial unfolding of H8 helix up to ASN337. (B-C) show the closed and open states of the midplane cavity and the hydrophobic gates. (D) Distinct 2D distributions of the H8 helicity and the cytoplasmic H8-H10 distance quantify partial unfolding of H8 and lateral opening at the cytoplasmic side under (+)PS-scrambling condition. (E) Distances between transmembrane helices H7, H8, and H10 at the bilayer midplane quantify lateral opening of the central cavity under the (+)PS-scrambling condition. (F) Minimum distances between hydrophobic-gate residues in the central cavity.

We have identified central hydrophobic gates at the midplane of the inner groove, formed by TYR278(H7):PHE344(H8) and TYR281(H7):TYR418(H9) (***Figure 3*B**). In (–)PS-scrambling simulations, minimum distances between these residue pairs indicate a stable closed-gate configuration, although the starting cryoEM-derived structure of hSERINC3 exhibits a conformational switch for PHE344, oriented away from the inner cavity (**Figure S5**). Under (+)PS-scrambling condition, both the hydrophobic gates formed between H7-H8 and H7-H9 widened by ∼ 5 Å (***Figure 3*F**, **Figure S1** C,F). Further, another hydrophobic gate has been identified near the cytoplasmic entrance, formed by PHE182(H4):TYR350(H8) residue pair, that also widened by ∼ 5 Å under (+)PS-scrambling condition (***Figure 3*B**, **Figure S2**).

### Pathway-dependent alterations of hSERINC3 central cavity without alternating access

In the classical “alternate access” model of lipid transporters, helical bundles undergo rigid-body movements around the central substrate-binding cavity, alternatively exposing it to the EC or IC side of the membrane. In hSERINC3, the charged residue pair E188(H4):R354(H8) in the IC leaflet remains separated by at least 1.5 nm throughout the (+)PS-scrambling simulations, indicating a stable inner-open state during lipid scrambling (**Figure *4*A**). Conversely, the separation distance between the polar residue pair S285(H7):N337(H8) in the EC leaflet remains below 1 nm (mostly), confirming an outer-closed state. These observations indicate that hSERINC3 does not follow an alternating access mechanism for PS scrambling (**Figure *4*A**, **Figure S4**A).

**Figure 4:**
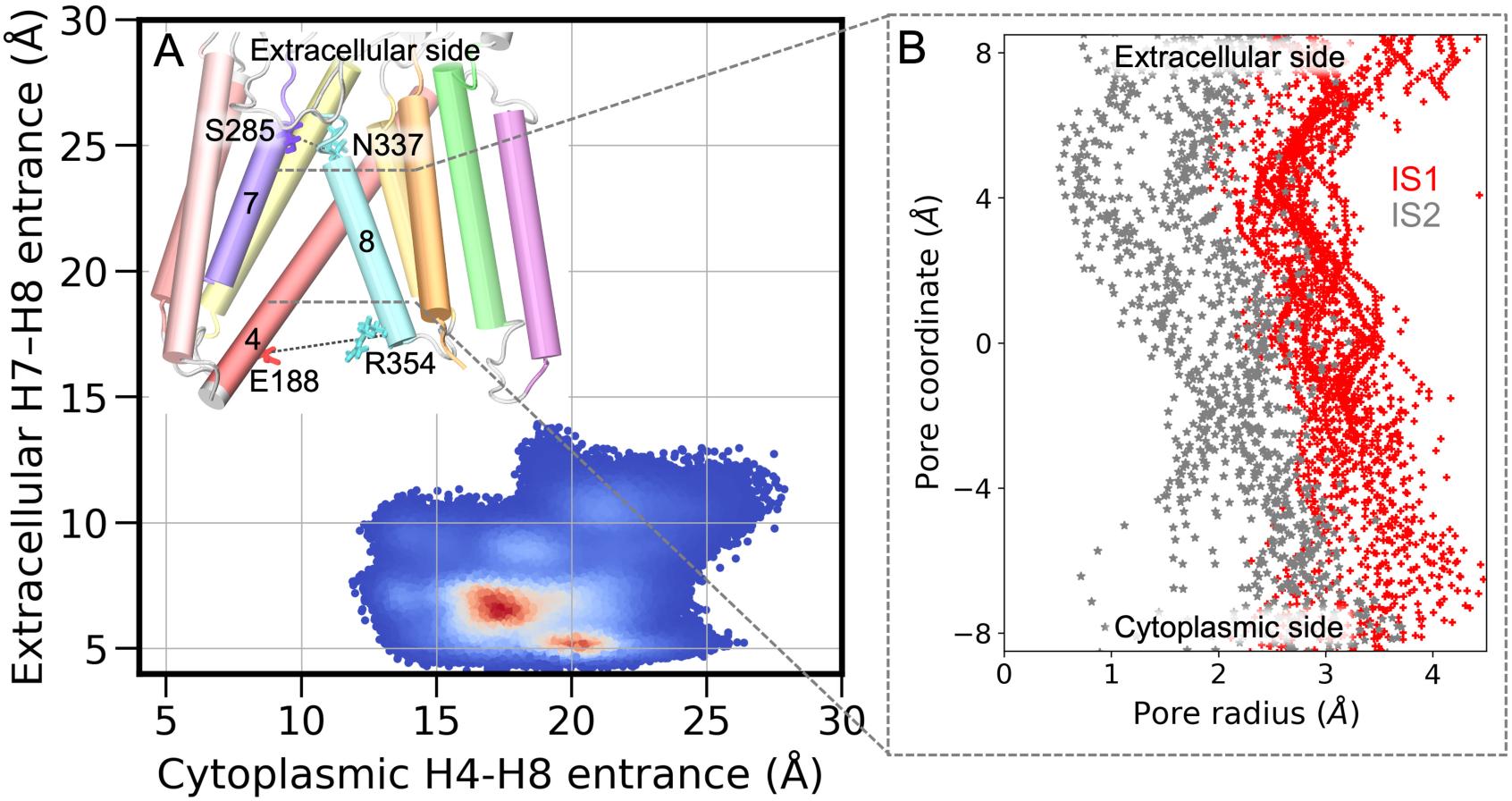
Dynamics of asymmetric cytoplasmic and extracellular lateral openings under the (+)PS-scrambling condition and pathway-dependent altered central cavity structure. (A) Distances between transmembrane helices lining the cytoplasmic and extracellular entrances. Calculations used the Cα atoms of charged and polar residues, highlighted in the hSERINC3 structure. Red regions indicate the most populated regions in the 2D space. (B) The pore radius profile along the axis perpendicular to the membrane, i.e., channel coordinate, for the hSERINC3 structures in the two intermediate metastable states, IS1 (red lines) and IS2 (gray lines). The extracellular and cytoplasmic ends of the pore are indicated.

We further examined the dynamics of inner cavity opening for different pathways of lipid scrambling. In the intermediate metastable state, IS1 of pathway1, where the lipid headgroup traverses the inner cavity, the pore is widely open compared to the IS2 conformational state of pathway2 (**Figure *4*B**), where the lipid headgroup interacts with protein surface residues. These results provide direct evidence for a closed-groove, non-canonical lipid scrambling mechanism.

### Wider lateral EC opening and the interactions between central cavity polar residues and PS headgroup stabilize the intermediate state of lipid scrambling

For different classes of scramblases following a credit-card mechanism, experimental studies reveal the spectrum of structural conformations adopted by the groove. Here, we investigate contributions of different conformations of hSERINC3 in stabilizing the intermediate state of lipid scrambling along pathway1. The FES, shown in **Figure 2**, suggests that at IS1, the PS lipid headgroup remains at the center of the protein inner groove (y-distance ∼ 0 nm, z-distance ∼ 0 nm). The cryo-EM derived hSERINC3 structure contains a narrow EC vestibule due to a constriction formed by the polar interaction network between H7, H8, and H9. Compared to the overall (+)PS-scrambling condition and cryoEM density map-derived structure, the EC end of hSERINC3 in the IS1 state exhibits a wider opening, still narrower than the IC end (**Figure *5*B**, **Figure S4**B).

**Figure 5:**
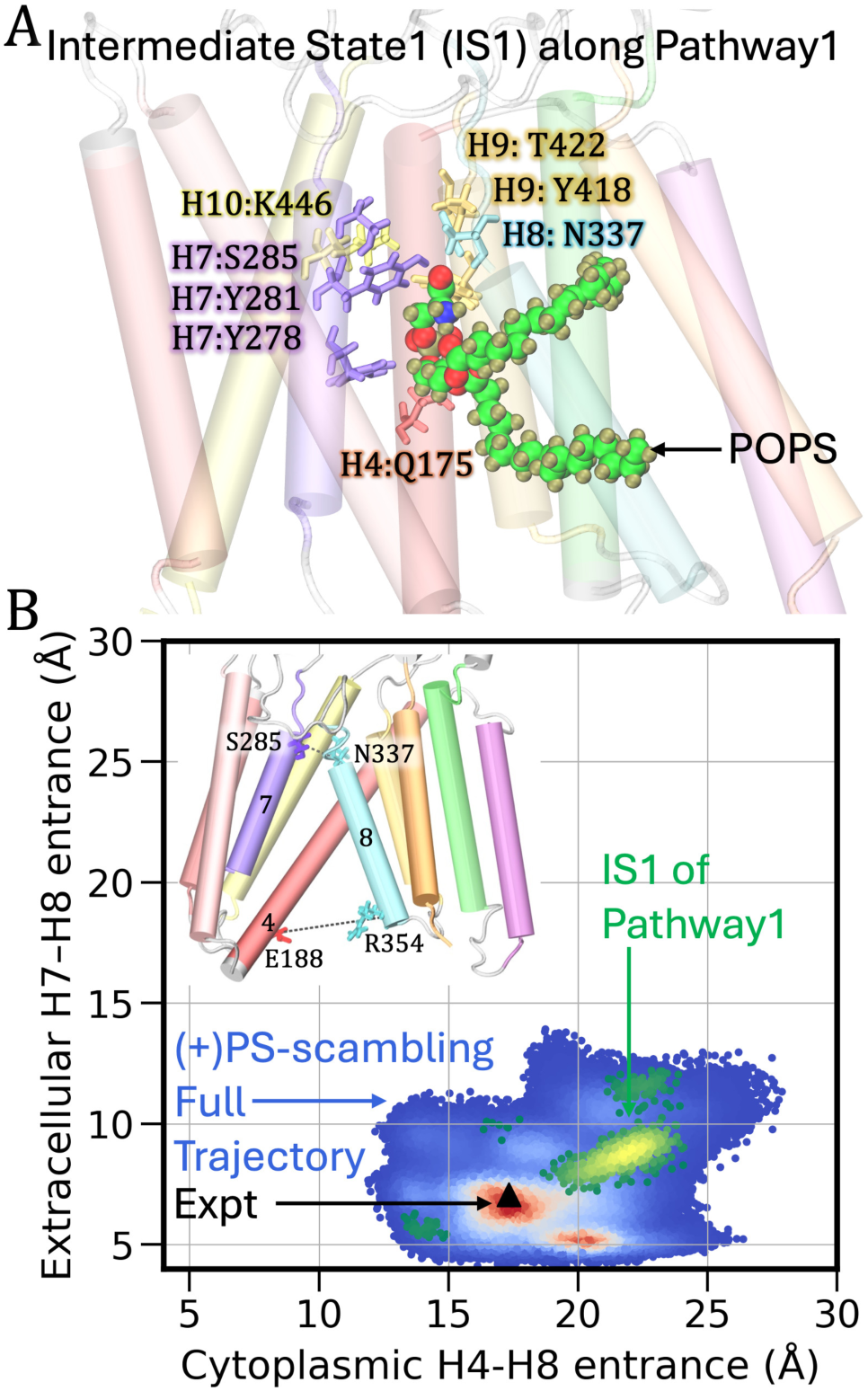
Specific interactions of hSERINC3 residues with the PS lipid headgroup and hSERINC3 conformational change stabilizing the intermediate state (IS1) of inner-groove lipid scrambling via the credit card mechanism. (A) hSERINC3 residues interacting with the PS lipid headgroup being translocated through the inner groove. N337(H8) in the unfolded conformation attains orientational flexibility to interact with the PS headgroup. (B) Characterization of the asymmetric cytoplasmic and extracellular openings in the IS1 state suggests that IS1 intermediate state is stabilized by the wider EC opening, still narrower than the IC opening. Yellow and red regions indicate the most populated regions in the 2D space for the IS1 state and the full (+)PS-scrambling trajectory, respectively. The black triangle indicates the corresponding value from the experimental cryo-EM structure.

A detailed analysis of protein-lipid contact frequency on the (+)PS-scrambling trajectories reveals crucial hSERINC3 residues that specifically interact with lipid headgroup at the IS1 state and facilitate PS lipid scrambling (see contact frequency data in **Figure S3** for different trajectories). The negatively-charged headgroup interacts with a ring of polar amino acid residues, formed by ASN337 of H8, GLN175 of H4, TYR278, TYR281, S285 of H7, TYR418, THR422 of H9 and LYS446 of H10 (**Figure *5*A**). In the (-)PS-scrambling trajectories, H8 remains in a helical form upto ASP334, and the orientation of the ASN337 residue in H8 remains away from the inner groove. Our (+)PS-scrambling simulations reveal that the partial unfolding of the H8 helix, discussed in the last section, facilitates the ASN337:PSheadgroup interactions.

## Discussion

By utilizing extensive AAMD with the TTMetaD approach, our study elucidates the molecular mechanism of PS scrambling by hSERINC3, revealing both a canonical “in-the-groove” pathway and a non-canonical alternate pathway through the surface of the protein. Scramblases are known to adopt different mechanisms depending on the lipid–protein interactions and protein conformational dynamics during the scrambling. Structurally, hSERINC3 resembles non-ATP dependent scramblases such as archaeal PfMATE, bacterial MurJ, and LtaA, all of which consist of two α-helical bundles connected by a long, tilted cross-member helix analogous to H4 in hSERINC3^26^. For MurJ and LtaA, experimental and computational studies have demonstrated an “alternating-access” mechanism, in which protein conformational switching between inward-open and outward-open states enables a “trap-and-flip” mode of lipid translocation, and in which the entire lipid molecule remains enclosed within the protein cavity during scrambling ^39,40^. In contrast, our simulations indicate that hSERINC3 does not undergo alternating-access conformational changes, and PS scrambling occurs via a “credit card” mechanism^13, 22^ in which lipid tails remain exposed to the membrane hydrophobic domain while the headgroup traverses through the inner hydrophilic cavity of hSERINC3. These findings establish hSERINC3 as a structurally similar but mechanistically distinct scramblase (compared with MurJ and LtaA).

For the first time, to our knowledge, our enhanced MD sampling simulations captured multi-pathway protein-mediated lipid scrambling and spontaneous close-to-open conformational transition of protein groove during lipid translocation(**Figure 2**). In pathway1, the PS headgroup scrambles through the central cavity of the protein. Remarkably, our simulations also captured a closed-groove scrambling mode in which the PS headgroup interacts with the residues of the hSERINC3 surface formed by NTD of H4, H6, and H10 (**Figure 2A**, **Figure 4B**). Such a mechanism has been hypothesized from indirect experimental evidence ^21^; for example, (i) the ability of afTMEM16 to scramble PEGylated lipids too large for the open groove^41^, (ii) failure to resolve WT TMEM16F scramblase structures with sufficiently wide open groove to accommodate lipids even under Ca^2+^-activated conditions^42, 43^. Recently, Grabe and coworkers attempted to unravel the alternate route of scrambling using the Martini coarse-grained models of a large number of scramblase structures in both open (Ca^2+^-bound) and closed (Ca^2+^-free) conformations ^21^. In our all-atom (+)PS-scrambling simulations, using a closed-groove initial state and without presuming the mechanism of lipid transport, the simulation sampled PS lipid scrambling in both in-the-groove and out-of-groove mechanisms, and the inner cavity opened up in an “unsupervised” way as the chosen collective variables, on which the bias potential was applied, do not dictate protein conformations. The FES contains combined energetic contributions of both protein-lipid interactions and protein conformational changes, which act synergistically during scrambling.

Our simulation method has allowed us to investigate atomistic details of protein conformational change during PS scrambling. We observed partial unfolding of the H8-NTD upto ASN337 under (+)PS-scrambling condition (**Figure 3D**, **Figure S1** A,D). For other ions and substrate transporters, the functional role of conformational transitions of transmembrane α-helices has been recognized through experiments ^44^ ^45^. For P4-ATPase, ATP8A (flippase), mutagenesis experiments have established the role of the transmembrane ASN residue in PS recognition and facilitating the flipping by stabilizing protein-PS headgroup interaction ^46^. Our study reveals that the helix-to-coil transition of ASN337 renders orientational flexibility of this polar residue at the inner cavity to interact with the PS headgroup and stabilizes the intermediate state (IS1) of the credit-card mechanism (**Figure 5A**). Intriguingly, the conformational transition of hSERINC3 in our (+)PS-scrambling simulations is consistent with the ill-defined density of H8 in the cryoEM density map of WT hSERINC3 (**Figure 3A**) ^26^, which not only helps to validate our simulation method, but also establishes the role of transmembrane ASN337 and conformational change of H8 in hSERINC3-mediated PS scrambling. Similar to hSERINC3, the AlphaFold model^47^ of hSERINC5 also contains an H8 ASN residue close to the central cavity (**Figure S10**). Interestingly, hSERINC2, which does not display restriction activity, does not have an H8 ASN residue.

An essential element in the credit-card mechanism of lipid scrambling is the opening of the inner hydrophilic groove to the membrane environment, and scramblases use different mechanisms of stabilizing the membrane-exposed conformations. For the (+)PS-scrambling condition, the distribution of separation distances of α-helices in the mid-plane of the bilayer, especially between H8-H7 and H8-H10, suggests a wide range of hSERINC3 inner groove structures regulates PS scrambling through different pathways (**Figure 3E**, **Figure S1** B,E, **Figure *4*B**). Interestingly, we identified hydrophobic gates of hSERINC3, formed by TYR and PHE residues of H4, H7, H8 and H9 (**Figure 3F**, **Figure S1** C,F, **Figure S2**). These gates remained closed in the unbiased AAMD simulation with cryo-EM derived structure of hSERINC3 under (-)PS-scrambling condition; however, a wide range of conformational states were observed under (+)PS-scrambling condition. The functional role of hydrophobic gates in regulating membrane transport of ions and lipids is well-recognized, and several mechanisms of gating have been proposed for different classes of transporters: an extracellular gate in TMEM16 scramblase ^48^, whereas a central activation gate in TMEM16F scramblase^49^. The ATP-dependent, flippase P4-ATPase^50^ also has a central activation gate. Instead of uncontrolled lipid translocation, the gates mediate regulated, signal-dependent lipid movement. All these proteins use hydrophobic gating and translocate lipids with a credit-card mechanism. On the other hand, lipid II flippase MurJ, which bears a structural resemblance with hSERINC3, translocates lipids by a “trap-and-flip” mechanism and exhibits an “alternating access” mechanism by large-scale protein conformational change, contains periplasmic and cytoplasmic gates formed by polar residues ^40^. In contrast, hSERINC3 contains both cytoplasmic and central hydrophobic gates. TYR350(H8):PHE182(H4) forms the hydrophobic gate near the cytoplasmic entrance of the protein groove, whereas TYR278(H7):PHE344(H8) and TYR281(H7):TYR418(H9) form the central gates near the inner hydrophilic groove of the protein. Intriguingly, the AlphaFold model^47^ of hSERINC5 also contains similar TYR residues in the central cavity and cytoplasmic entrance domains (**Figure S10**), which are not present in hSERINC2. While understanding the specific functions of these gates is beyond the scope of this work, we can speculate that the central hydrophobic gate can regulate lipid headgroup specificity.

Structural characterization of different classes of lipid transporters has revealed the mechanism of lipid translocation following an alternate access mechanism, facilitated by the large-scale protein conformational transition between inward-open and outward-open states. Also, deviation from the canonical alternate-access model has been proposed for ABC transporters, PglK-catalysed lipid-linked oligosaccharide flipping that follows a “outward-only” mechanism ^17^. The experimental structure of hSERINC3 contains a narrow transmembrane aqueous channel (0.1 nm) that remained stable in multiple microsecond-long unbiased AAMD simulations, which did not sample any PS flipping events. The analysis of dynamic opening/closing of cytoplasmic and extracellular entrance under the (+)PS-scrambling condition displayed wider lateral opening on the cytoplasmic side of the membrane (>1.5 nm) with a closed extracellular entrance (<0.8 nm) (**Figure 4**). This is consistent with other replicas of the TTMetaD simulations (**Figure S4**A) and suggests that the central cavity of hSERINC3 is not accessible alternatively following the classical alternating access model. Our simulations further allowed us to perform structural characterization of the intermediate state of the credit card mechanism lipid scrambling (**Figure 5**, **Figure S4**B). For different scramblases, following the credit card mechanism, structural biology experiments have revealed several conformational states such as “membrane-exposed”, “lipid-conductive” etc. ^51^, however, their contributions are not yet fully understood. Our simulation method is able to determine the conformational state of hSERINC3, stabilizing the intermediate state of inner-groove PS scrambling. The EC end of hSERINC3 in the IS1 state of pathway1 exhibits a wider opening, compared to the overall (+)PS-scrambling condition and the cryoEM density map-derived structure (**Figure 5B**). However, the IC opening at this intermediate state (>2 nm) is still significantly wider than the EC opening (<1 nm). Additionally, our simulation method allowed us to determine the specific hSERINC3:PS headgroup interactions stabilizing the intermediate states (**Figure 5A**, **Figure S3**).

The simulation method presented here explored pathways and energetics of a PS scrambling mediated by hSERINC3. However, a fluorescent, proteoliposome flipping assay demonstrated that hSERINC3 can also scramble PC and PE lipids^26^. Our current computational method does not allow the investigation of the correlation between multiple lipid flip-flops. Although we have observed correlated movement of adjacent cholesterol molecules during PS lipid scrambling, further study is required to understand the correlation between hSERINC3-mediated PS lipid scrambling and cholesterol flip-flop motion in detail. Additionally, whether any specific ion facilitates the PS scrambling by hSERINC3 is not known. The present study aimed to understand the hSERINC3-PS specific interactions and hSERINC3 conformational change during PS lipid scrambling.

In summary, our study suggests that (1) PS can be scrambled by hSERINC3 through multiple pathways (**Figure *2***), (2) PS scrambling is facilitated by partial unfolding of H8, opening of hydrophobic central gates and central cavity by rearrangement of transmembrane α-helices (**Figure *3***), (3) Wider EC lateral opening by the relaxation of EC constrictions (formed by polar residue network between H7, H8 and H9) and the interactions between polar residues in the central cavity with PS headgroup stabilize the intermediate metastable state of inner-groove PS scrambling that follow a credit-card model (**Figure 5**), (4) The central cavity is not accessible alternatively from EC and IC ends as described by a classical “alternating access” model and adopts different conformations in the intermediate states of pathway1 and pathway2 of PS scrambling (**Figure *4***), (5) PS lipid scrambling perturbs conformations of helical bundles of hSERINC3 significantly during scrambling, which is not consistent with the rigid body movement of two α-helical bundles, as accepted for a typical alternating access model for membrane transport (**Figure S7**). Overall, the mechanistic elements described here reveal that (1) the PS lipid scrambling mechanism of hSERINC3 is different than the other non-ATP-dependent transporters having similar architecture, thereby expanding the paradigm of lipid scrambling and (2) the key residues of hSERINC3 for PS scrambling, namely N337 in H8 and a cluster of tyrosine residues in the central cavity that form the central gate and interact with the PS headgroup, are also present in the central cavity of hSERINC5 (AlphaFold model, **Figure S10**) that exhibits the maximum restriction activity against HIV-1. Although the mechanism of viral restriction by hSERINC5 and hSERINC3 – and the extent to which their lipid scrambling activity contributes to this function – remain unresolved, our present study elucidates the molecular-level mechanism of hSERINC3-mediated PS scrambling and identifies that the shared structural features between hSERINC3 and hSERINC5 play a crucial role in PS scrambling which are not present in hSERINC2 that lacks antiviral activity.

## Methods

### System and Simulation Setup

All AAMD simulations were performed using the structure of hSERINC3 derived from cryo-electron microscopy (cryo-EM) (PDB ID: 7RU6) ^26^. A fully hydrated bilayer around the protein was built and equilibrated using the CHARMM-GUI Membrane Builder and Quick-Solvator, respectively^52^. The membrane-embedded protein was charge-neutralized and solvated in 150 mM aqueous NaCl solution with initial box lengths of 12 nm × 12 nm × 10.5 nm to allow sufficient water layers to be present between the protein and its periodic image. Five calcium ions were added to each system to mimic physiologic calcium concentration. The total size of the systems was ∼150,000 atoms. The lipid bilayer model designed for this study was based on lipidomic analysis of HIV-1 particles^53^, and we have selected a complete asymmetric distribution of PS in the initial structure of the bilayer. The composition of the extracellular leaflet was 40% cholesterol, 15% phosphatidylcholine (PC), and 30% sphingomyelin (SM), and the cytoplasmic leaflet was 25% cholesterol, 20% PC, 20% phosphatidylethanolamine (PE), 30% phosphatidylserine (PS), and 8% phosphatidylinositol (PIP2). We have considered 8% PIP2, 30% PS in the inner leaflet, which is 3% and 12% of the total number of lipid molecules in the bilayer (Please note here that the outer leaflet contains a greater number of lipid molecules than the inner leaflet in an asymmetric membrane due to the difference in lipid packing).

We used the CHARMM36m force field ^54^ for protein and lipids, and CHARMM TIP3P for water ^55^. Simulations were performed using GROMACS 2020.4 MD software ^56^. Energy minimization was performed using the steepest descent algorithm until the maximum force was less than 1,000 kJ mol^−1^nm^−1^. After the energy minimization step, the equilibrium stage was divided into a 6-step protocol provided by CHARMM-GUI, consisting of two NVT steps and four NPT steps in sequence for a total of 26 ns. The previous harmonic positional restraints were removed, and the system was equilibrated in a constant NPT ensemble for an additional 200 ns. Next, 2 μs trajectories were generated for 4 replicas. Throughout this procedure, the temperature was kept constant at 310.15 K using the Nosé -Hoover thermostat with a 1.0 ps coupling constant ^57, 58^, and the pressure was set at 1 bar and controlled using the Parinello-Rahman barostat semi-isotropically due to the presence of the membrane. The compressibility factor was set at 4.5 ×10^-5^ bar^-1^ with a coupling time constant of 5.0 ps ^59^. Van der Waals interactions were computed using a force-switching function between 1.0 and 1.2 nm, while long-range electrostatics were evaluated using Particle Mesh Ewald(PME) ^60^ with a cutoff of 1.2 nm, and hydrogen bonds were constrained using the LINCS algorithm ^61^.

### Transition-tempered Metadynamics (TTMetaD) Simulations

The TTMetaD method was used to sample hSERINC3-mediated phosphatidylserine (PS) lipid scrambling events^30^. TTMetad converges rapidly and asymptotically without sacrificing exploration of configurational phase space in the early stages of simulations (**Figure S9**). Several efficient enhanced sampling methods (such as umbrella sampling, well-tempered metadynamics) were not adequate for this problem of interest. In addition to lipid flipping, there are other competitive processes, such as PS lipid lateral diffusion away from hSERINC3 protein and lipid desorption from the membrane. TTMetaD outperforms well-tempered metadynamics in terms of convergence, as it mainly explores the free energy surface around two defined basins and converges rapidly and asymptotically. For the lipid scrambling process, the two basin points are well-defined:

(1) the PS lipid headgroup at the inner leaflet and (2) the PS lipid headgroup at the outer leaflet. Two collective variables (CVs) for TTMetaD were used to characterize SERINC3-mediated PS lipid scrambling events: (1) z-distance of the center-of-mass of PS lipid headgroup from hSERINC3 center-of-mass (COM), which is the scrambling coordinate. z<0, z>0, and z∼0 are when the PS headgroup is at the inner leaflet, the outer leaflet, and close to the bilayer center, respectively. (2) y-distance of the center-of-mass of the PS lipid headgroup from the hSERINC3 COM, which varies depending on whether the PS lipid headgroup interacts with the protein surface or the central groove. We applied lower and upper harmonic walls to limit the phase space accessible to the system during the simulation. There is another unbiased CV, x-distance of the COM of the PS lipid headgroup from the bilayer center, which was carefully monitored for the proper exploration of phase space, instead of getting trapped close to the lower and upper harmonic walls.

Well-equilibrated protein-membrane initial structures after 2 μs AAMD simulations were used for TTMetaD simulations. All TTMetaD simulations were run using GROMACS version 2020.4 patched with PLUMED 2.7.0 open-source library ^62^. The trajectories are shown in **Figure S8**. The initial Gaussian hill height for TTMetaD simulations was 2.0 kJ/mol with a width of 0.2 nm for both distance CVs. The pace of hill deposition was every 500 timesteps or 1 ps. The TTMetaD bias factor was 11. In the production runs, the configuration of the system was saved every 10 ps interval. The free energy profile of protein-lipid interactions was estimated employing the reweighting algorithm, proposed by Tiwary *et al.* ^63^ in order to recover unbiased distributions.

### Analysis of the All-atom MD Simulations

Protein-lipid interactions were visualized using VMD 1.9.3 ^64^. Analyses of the trajectories were performed using GROMACS^56^, PLUMED^62^, MDAnalysis^65^, and VMD Tcl scripts^64^. To compute the properties of (-)PS-scrambling simulations (**Figures 1B, 3D–F** in the main text; **Figures S1, S2, and S6** in the Supporting Information), the final 700 ns of 4 unbiased AAMD trajectories were concatenated. The MEPSA Python package was used to compute minimum energy pathways in **Figure *2*A**^66^. The helicity of H8 (residues 334-354) (in **Figure *3*D**) was computed using PLUMED^62^ Colvar ALPHARMSD, which measures the number of all sets of contiguous six residues in alpha helical form. Therefore, a value of 16.0 for this parameter signifies that all 21 residues in the segment residue 334-354 are in alpha helical conformation. For the calculation of cytoplasmic H8-H10 distance (in **Figure *3*D**), we considered the distance between the C*α* atoms of R354 (H8) and A463(H10). Midplane H8-H7 and H8-H10 distances (in **Figure *3*E**) were computed using the COM of residues in those corresponding helices within 0.5 nm above or below the bilayer middle plane. For the analysis of the hydrophobic gate, we considered the minimum distances between pairs of residues. The analysis of the pore radius (**Figure *4*B**) was performed using the HOLE program^67^ implemented in MDAnalysis^65^. We further quantified the local twist and bend angles of H4 helix residues (162-182) using the HELANAL module^68^ of MDAnalysis^65^ (**Figure S6**). Dimensionality reduction with time-lagged independent component analysis (tICA) ^38^ was performed using MDTraj^69^ and MSMBuilder^70^ (**Figure S7**).

## Supporting information

Supporting Information - Mechanism of Phosphatidylserine Lipid Scrambling by Human SERINC3, an HIV-1 Restriction Factor

## Data availability

All data needed to evaluate the conclusions in the paper are present in the paper and/or the Supplementary Materials. Simulation model and input files are uploaded in Zenodo: 10.5281/zenodo.17567096.

## Acknowledgments

This research was supported by the Frost Institute for Chemistry and Molecular Science at the University of Miami to MY. This research was also supported in part by the National Institute of Allergy and Infectious Diseases (NIAID) of the National Institutes of Health (NIH) by grant U54 AI170855 for the Behavior of HIV in Viral Environments (B-HIVE) Center (to G. A. V.). The content is solely the responsibility of the authors and does not necessarily represent the official views of the National Institutes of Health. The authors acknowledge computational resources provided by the University of Chicago Research Computing Center (RCC), the NIH-funded Beagle-3 computer (NIH award 1S10OD028655-01), and the Frontera supercomputer at the Texas Advanced Computer Center funded by the National Science Foundation (OAC-1818253).

## Author Contributions

M.Y., G.A.V. and P.B. conceptualized the research. P.B. prepared simulation models, designed the method and performed simulations, analyzed the data, prepared the figures, and wrote the original draft. M.Y. and G.A.V. supervised the research, writing process, and contributed to the discussions to finalize the manuscript.

