## Supporting Information - Mechanism of Phosphatidylserine Lipid Scrambling by Human SERINC3, an HIV-1 Restriction Factor for "Mechanism of Phosphatidylserine Lipid Scrambling by Human SERINC3, an HIV-1 Restriction Factor"

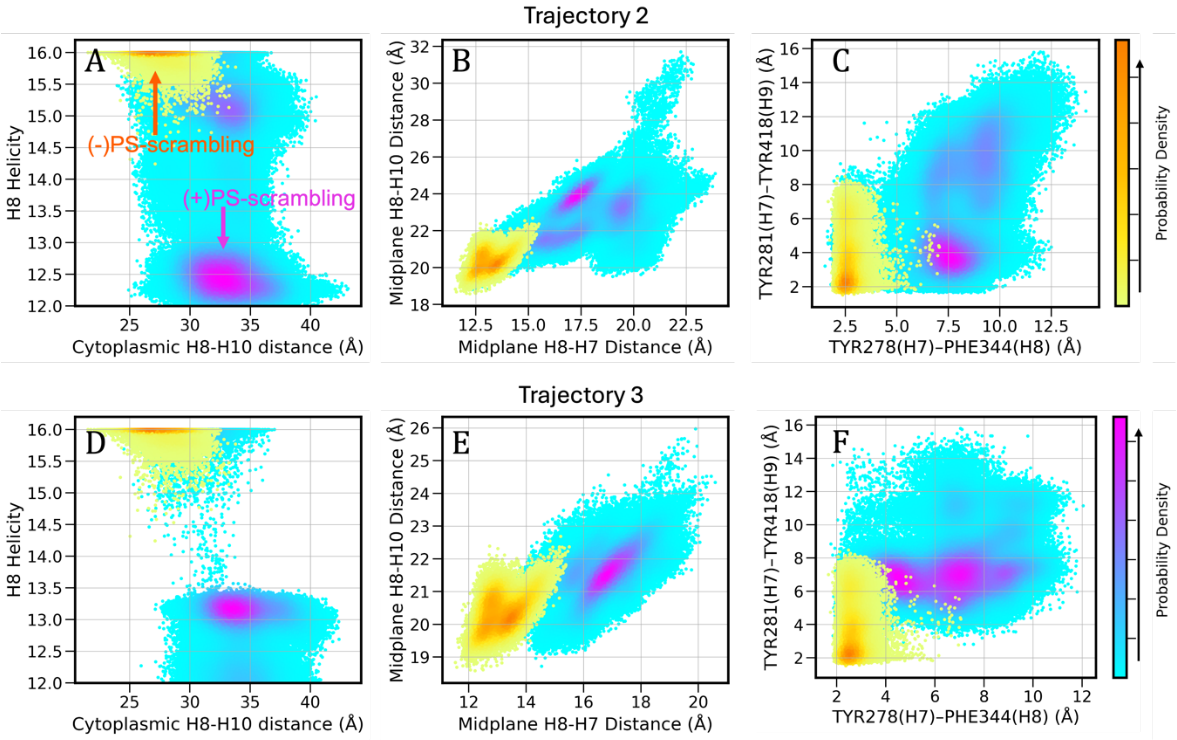


**Figure S1: hSERINC3 conformational dynamics mediating PS scrambling.** *(A, D)* Distinct 2D distributions of H8 helicity and cytoplasmic H8-H10 (R354-A363) distances sampled in (-)PS-scrambling and (+)PS-scrambling simulations. (B,E) Analysis of distances between TM helices (H7, H8, H10) at the bilayer middle plane and (C,F) the distances between TYR and PHE residues forming the hydrophobic gates at the central cavity. It is important to note that the difference in the distance distributions in different trajectories reflects the conformational variability, especially the dependence of hSERINC3 conformations on the local membrane environment in different trajectories.

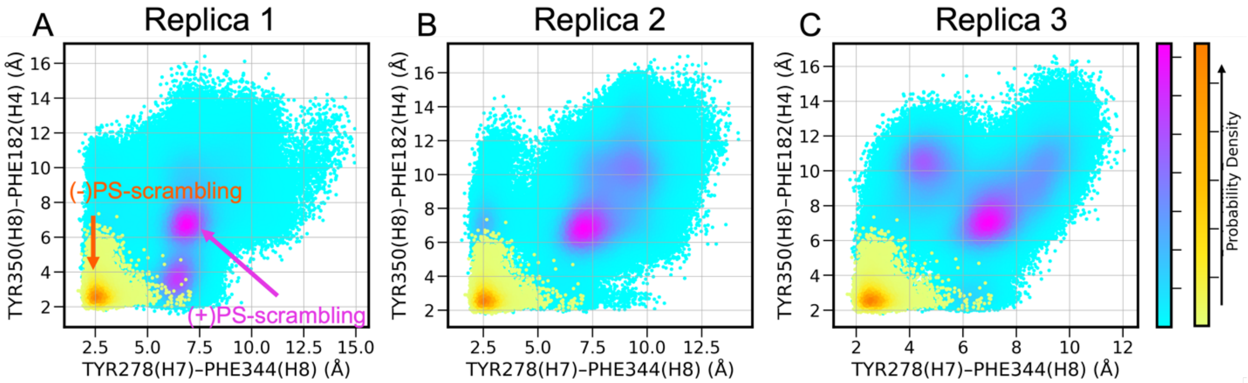


**Figure S2: Opening of hSERINC3 cytoplasmic and central hydrophobic gates under (+)PS-scrambling condition.** TYR350(H8):PHE182(H4) forms the hydrophobic gate near the cytoplasmic entrance of the protein groove. TYR278(H7):PHE344(H8) forms the central gate near the inner hydrophilic groove of the protein. (A-C) Probability distribution of minimum distances between the pair of residues forming the gates in different TTMetaD trajectories. Like Figure S1, the difference in the distance distributions in different trajectories reflects the conformational variability, especially the dependence of hSERINC3 conformations on the local membrane environment in different trajectories.

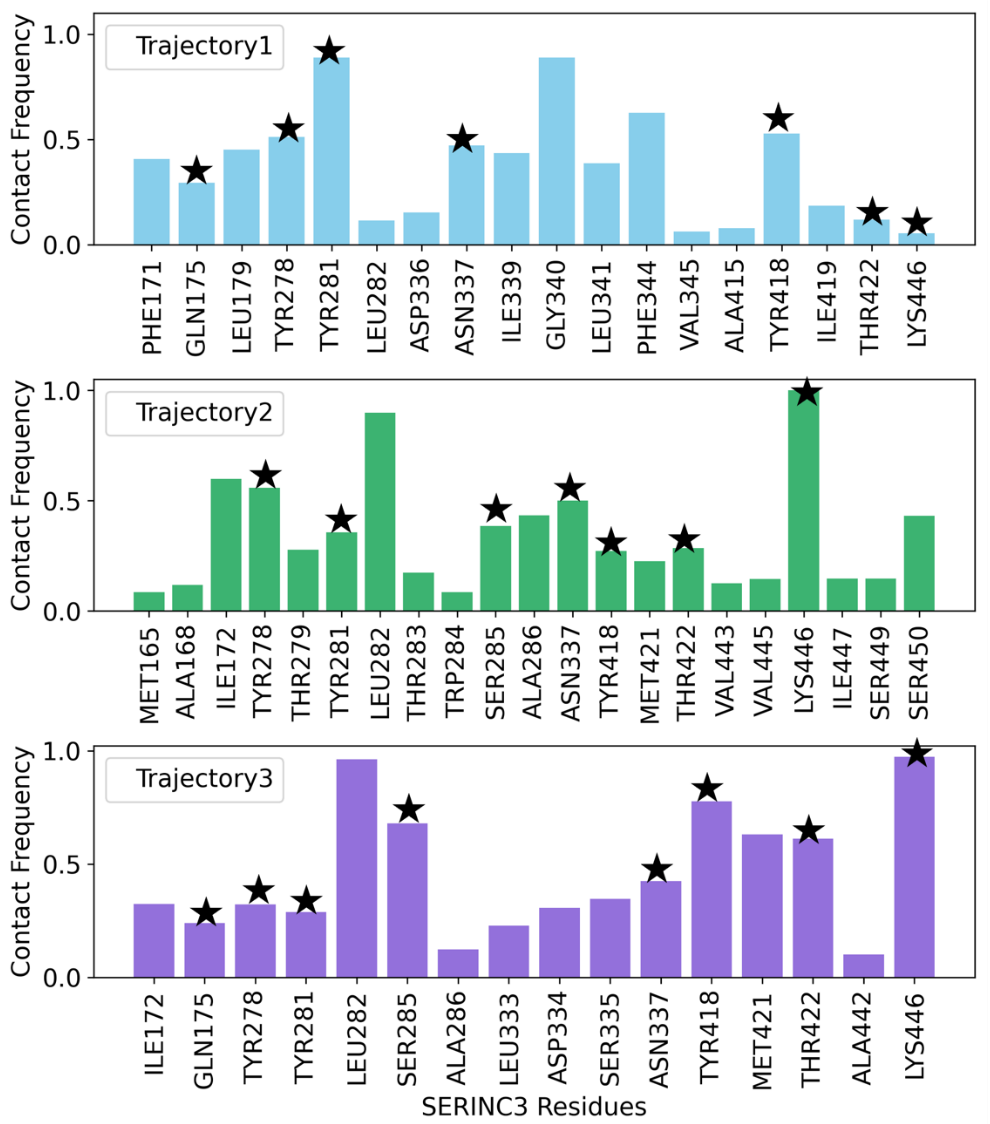


**Figure S3: Specific interactions between hSERINC3 and PS lipid headgroup at the intermediate state 1 (IS1) of PS scrambling.** PS lipid headgroup is considered to be in contact with hSERINC3 residues if any of their atoms are within a cutoff distance of 0.35 nm from any atom of that amino acid residue. The common contact residues in three trajectories are highlighted with stars. The simulation snapshot of the protein inner cavity with these residues is shown in **Figure 5A** of the main text.

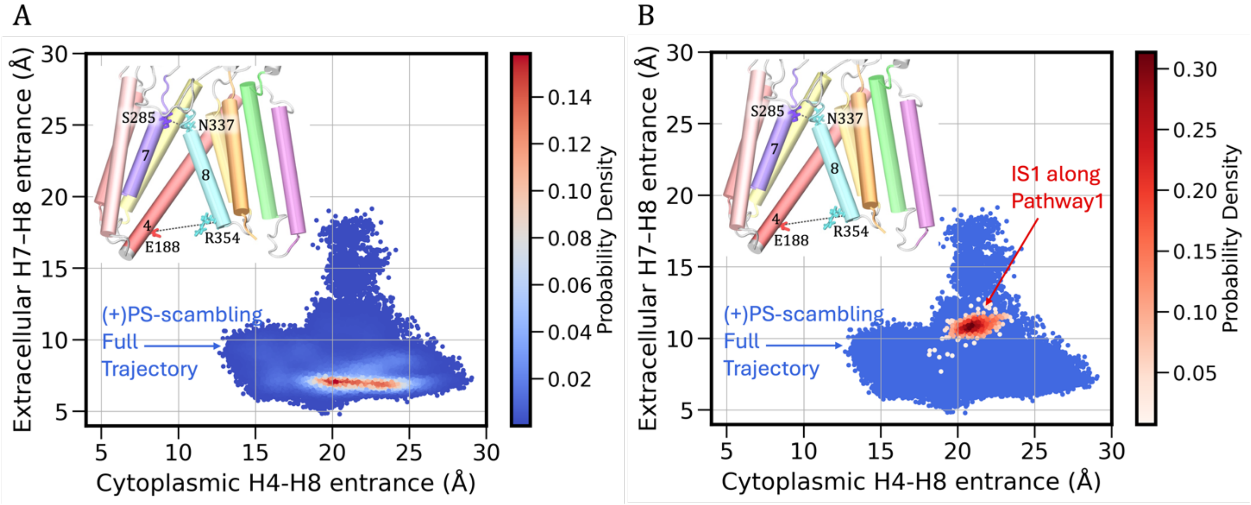


**Figure S4: Dynamics of asymmetric lateral EC and IC openings in replica2.** Calculations used the distance between Cα atoms of charged and polar residues, highlighted in the hSERINC3 structures. (A) Full trajectory of (+)PS-scrambling condition, (B) the intermediate states of inner-groove lipid scrambling were considered for the calculations. Only the scatter plot distance distribution of the full trajectory is shown in blue in (B) with the data for IS1 shown as a density map, where the dark brown region indicates the most populated IS1 states. Cytoplasmic and extracellular opening in the IS1 intermediate metastable state suggests wider EC opening, still narrower than IC opening, stabilizes this intermediate state, as also happened in replica1 shown in **Figure 5** of the main text.

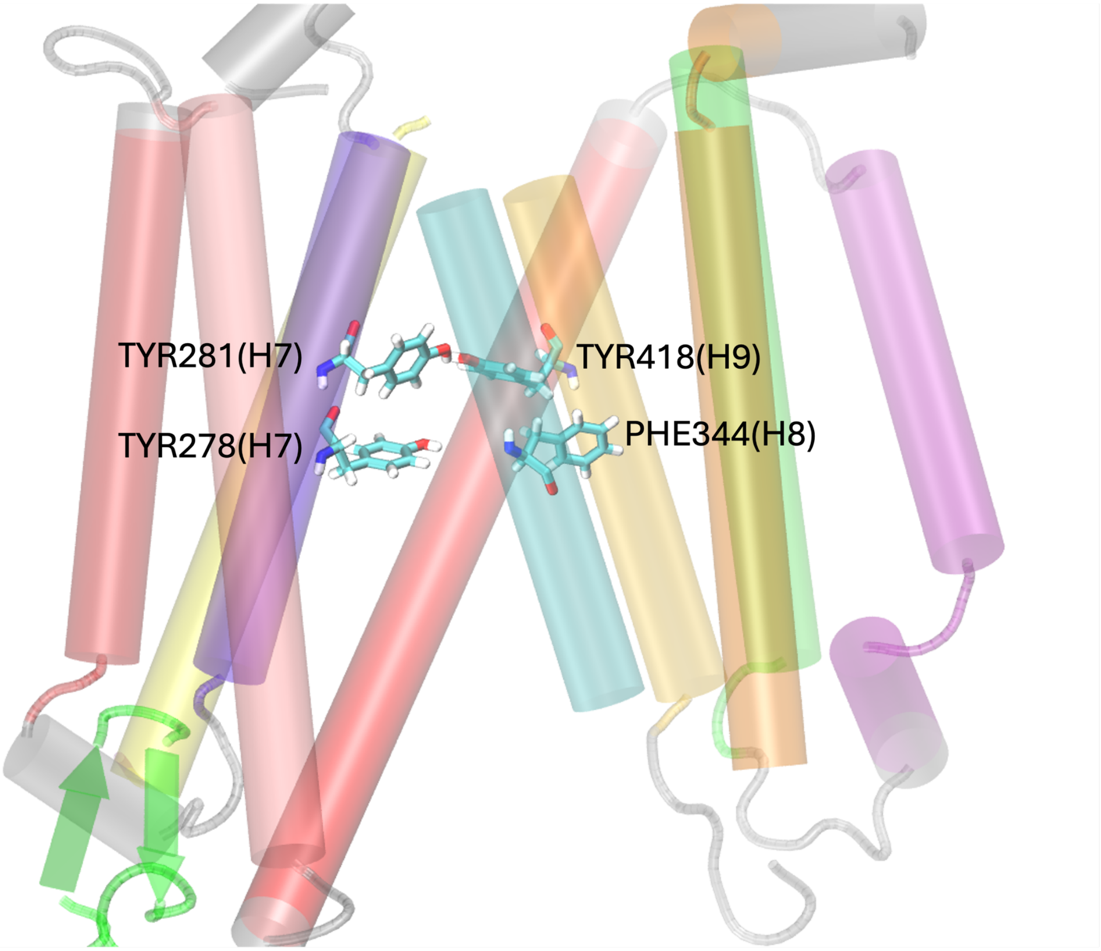


**Figure S5: Central gate-forming residues in the WT hSERINC3 cryo-EM structure with average resolution 4.2 Å (PDB ID: 7RU6).** In this structure, the PHE344 residue is turned away from the inner cavity and does not participate in hydrophobic gate formation. However, unbiased MD simulations initiated with this structure sampled configurations in which PHE344 spontaneously rotated toward the inner cavity, forming a closed central gate (see **Figure 3B, F** of the main text).

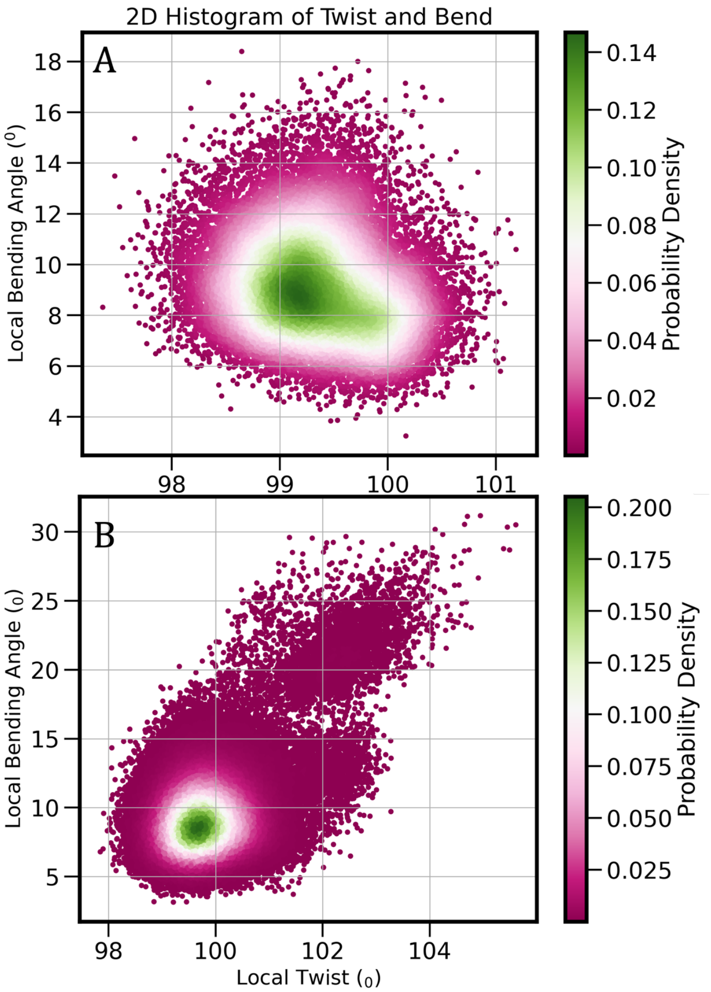


**Figure S6: Variations in helical parameters.** Local twist and local bending angles for crossmember H4 helix (residues 162-182) under (A) (-)PS-scrambling, (B) (+)PS-scrambling conditions.

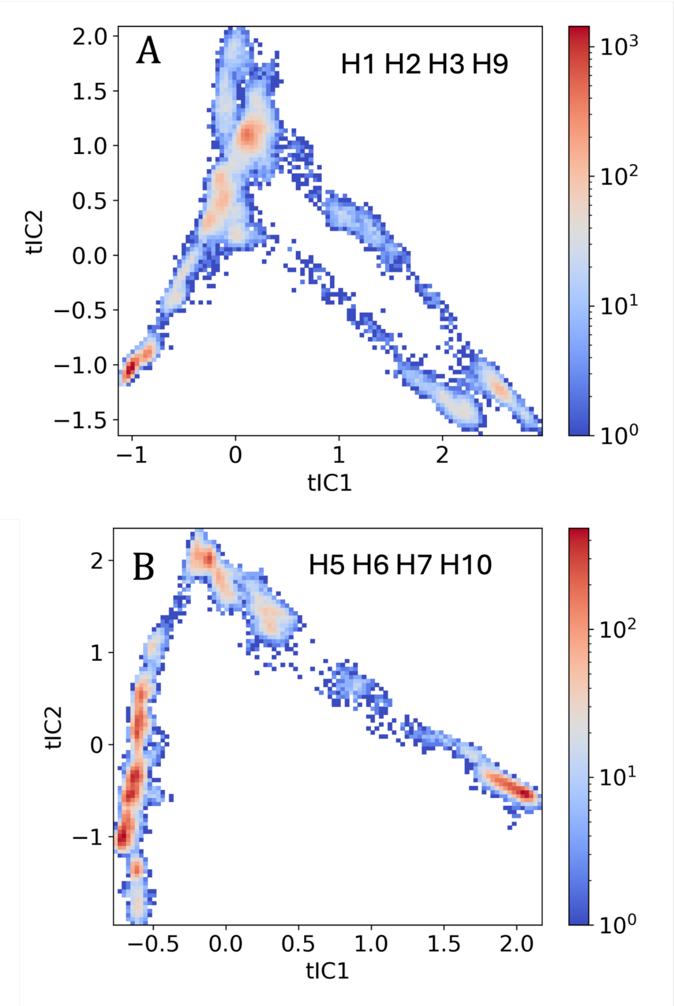


**Figure S7: Non-rigid movement of helical bundles during lipid scrambling captured by 2D tICA space spanned by the first two vectors.** We chose the variables, pairwise C$\alpha$-C$\alpha$ distance between residues of the helical bundles in the transmembrane domain, to compute the covariance matrix. tICA heatmaps for (A) H1-H2-H3-H9 and (B) H5-H6-H7-H10 helical bundles suggest multiple conformational states of these helical bundles under (+)PS-scrambling conditions. Both the tilt angle changes and the COM distance change of individual helices in the bundles can contribute to this conformational variability, which provides direct evidence that hSERINC3 does not undergo rigid-body motion of helical bundles as described in the classical alternate access model.

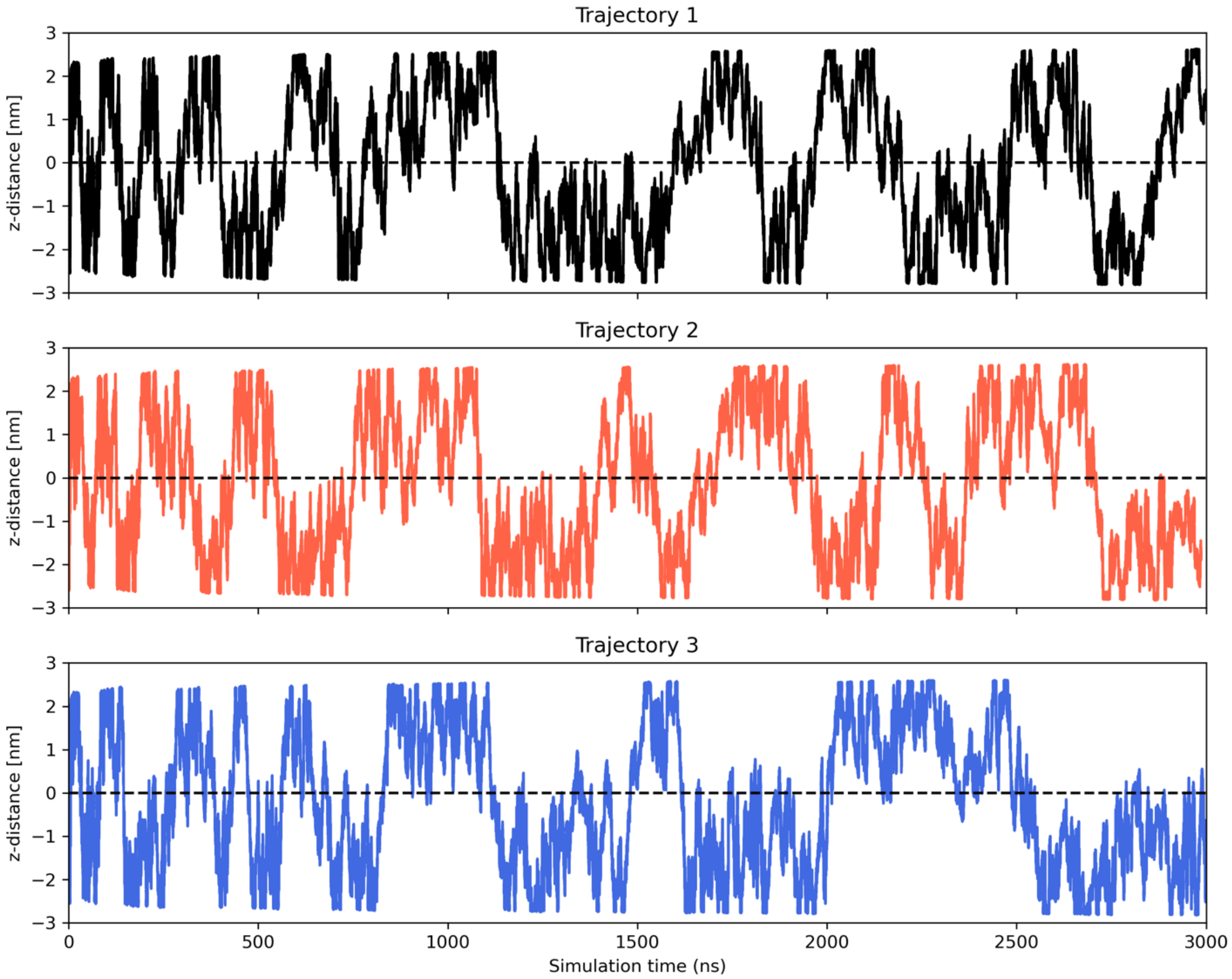


**Figure S8: The time evolution of the collective variable (CV) of TTMetaD trajectories**. The z-distance between the COM of the PS lipid headgroup and the hSERINC3 center-of-mass (COM), which is the scrambling coordinate. When the lipid headgroup is at the intracellular leaflet, the center of the bilayer, and the extracellular leaflet, the value of this CV is -ve, ~0, and +ve, respectively. Within 3 μs, trajectories sample ≥16 PS scrambling events. Please note that the timescale of this enhanced sampling simulation is defined differently compared to the unbiased MD simulation.

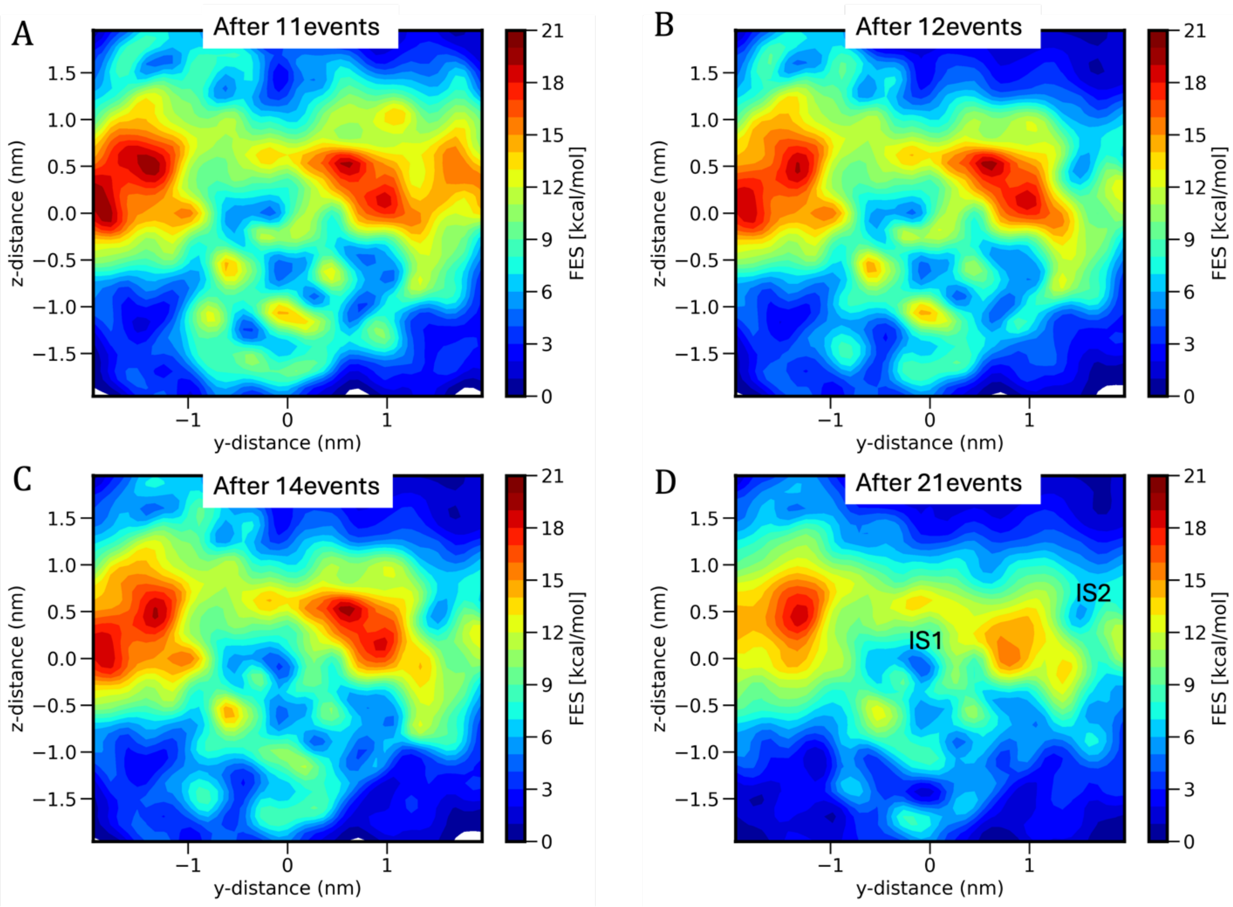


**Figure S9: Convergence of TTMetaD simulations.** Free energy surface of hSERINC3-PS headgroup interactions during PS scrambling as a function of two collective variables, z-distance and y-distance of the COM of PS headgroup from hSERINC3 COM.

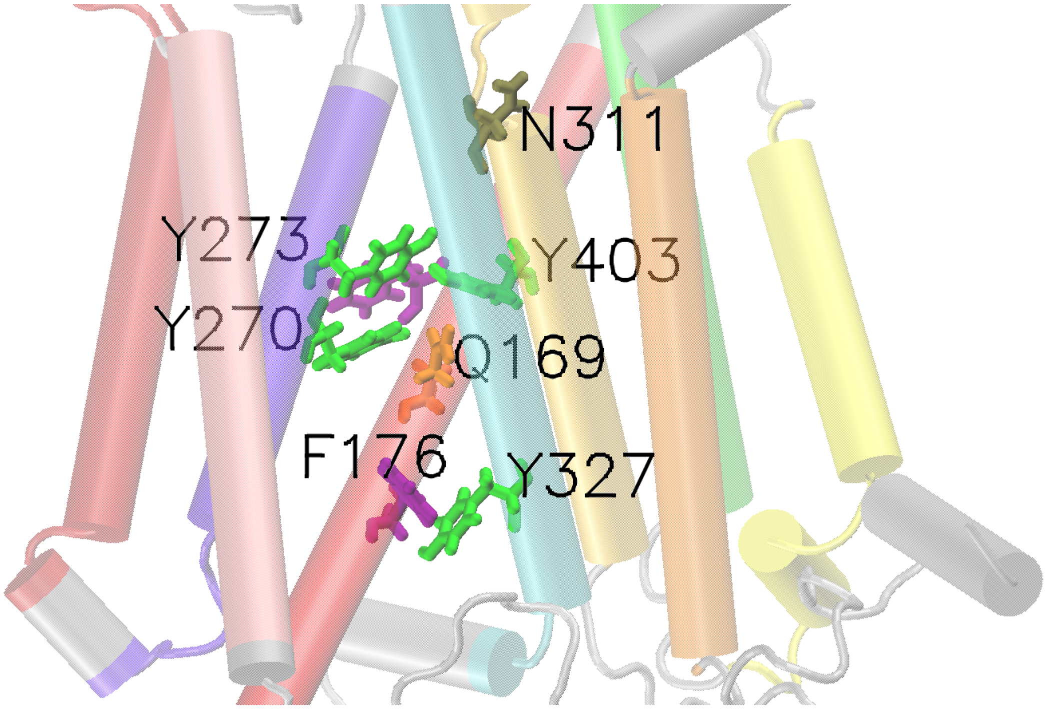


**Figure S10: Alphafold model of hSERINC5.** Residues analogous to those in hSERINC3, which are crucial for hSERINC3-mediated phosphatidylserine (PS) scrambling, are highlighted. These residues interact with the PS headgroup in the intermediate metastable state of inner-groove scrambling and contribute to the formation of the central and cytoplasmic gates.
